# Insight into Polyproline II Helical Bundle Stability and Folding from NMR Spectroscopic Characterization of the Snow Flea Antifreeze Protein Denatured State

**DOI:** 10.1101/2022.03.10.483783

**Authors:** Miguel Á. Treviño, María Redondo Moya, Rubén López Sánchez, David Pantoja-Uceda, Miguel Mompeán, Douglas V. Laurents

## Abstract

The use of PPII helices in protein design is currently hindered by limitations in our understanding of their conformational stability and folding. Recent studies of the snow flea antifreeze protein (sfAFP), a useful model system composed of six PPII helices, suggested that a low denatured state entropy contributes to folding thermodynamics. To get atomic level information on the conformational ensemble and entropy of the reduced denatured state of sfAFP, we have analyzed its chemical shifts and {^1^H}-^15^N relaxation parameters by NMR spectroscopy at three experimental conditions. No significant populations of preferred secondary structure were detected. The stiffening of certain N-terminal residues at neutral versus acidic pH leads us to suggest that favorable charge-charge interactions could bias the conformational ensemble to favor the formation of the two disulfide bonds during nascent folding. Despite a high content of flexible glycine residues, the mobility of the sfAFP denatured ensemble is similar for denatured α/β proteins both on fast ps/ns as well as slower μs/ms timescales. These results are in line with a conformational entropy in the denatured ensemble resembling that of typical proteins and suggest that new structures based on PPII helical bundles should be amenable to protein design.

## Introduction

Great advances have been made in recent years in protein design. Novel protein folds as well as new enzymes catalyzing reactions which are unknown in nature have been achieved (1). Nevertheless, almost all are based on combinations of a-helices or β-sheets or both, but largely ignore polyproline II (PPII) helices (2). In nature, PPII helices are common as they are the main component of collagen, the most abundant human protein, and most globular proteins contain at least one turn of PPII helix (3)(4)(5). Moreover, a small protein class, recently reviewed (6), which includes snow flea antifreeze proteins (sfAFP) (7), bacteriophage tail spike proteins (8), the enzyme acetophenone carboxylase (9), the universally conserved translation factor Obg (10) and the recently discovered anaplastic lymphoma kinase (11) consist of or contain glycine-rich PPII helical bundle domains. These glycine-rich PPII helical bundles can give rise to remarkably flat or honeycomb structures, which could find novel applications in protein design and biotech applications. Nevertheless, their use in protein design is hampered by a lack of understanding regarding the bases of their conformational stability.

Recently, we characterized the stability and dynamics of the *Hypergastrum harveyii* sfAFP by NMR spectroscopy and computational methods (12). This protein’s sequence and 3D structure, which was determined by X-ray crystallography (7) and consists of six long PPII helices connected by short turns, are shown in **Sup. Fig. 1**. The results showed that sfAFP is as well folded and rigid as globular proteins composed of a-helices or β-sheets or both despite a very high content of glycine residues. In fact, the composition of sfAFP is almost 50% glycine residues. The presence of two disulfide bonds and 28 weak Cα-H···O=C contribute significantly to the conformational stability of sfAFP (12) (13). However, glycine residues have very small side chains -just H- and can sample broad regions of the Ramachandran space. This means that the conformational entropy change for folding is expected to be much larger and unfavorable for glycine residues relative to the other eighteen common amino acid residues and proline. Nevertheless, the residue level dynamics and conformational entropy of the unfolded sfAFP are still unknown.

To address this issue, the goal of this study is to characterize the dynamics of unfolded sfAFP at the level of single residues using NMR spectroscopy. Previous NMR characterizations of unfolded proteins have made use of chemical denaturants, such as urea (14) (15). On the other hand, disulfide bond reduction is perhaps a more physiologically relevant way to denature proteins. In contrast to glycine residues, intact disulfide bonds act to stabilize proteins by limiting the conformational freedom of the denatured state (16). In this way, they decrease the unfavorable change in conformational entropy that accompanies folding. Using theory, it is possible to calculate the stabilizing contribution based on the number of residues within the loops closed by the disulfide bonds (17) and is 7.2 kcal/mol at 25.0 °C in the case of sfAFP. Considering that the overall conformational stability of sfAFP is significantly lower than 7.2 kcal/mol (12) (13), disulfide bond reduction is expected to denature this protein, as was observed by low/medium resolution spectroscopic methods (13). Here, we confirm by multidimentional NMR spectroscopy that sfAFP is indeed unfolded by reduction. The chief result of this study is that ps/ns and μs/ms backbone dynamics of sfAFP, as measured by ^1^H-^15^N based NMR relaxation measurements, retains a moderate rigidity akin to other disordered or unfolded proteins with a much lower glycine content.

## Materials and Methods

sfAFP isotopically labeled with ^13^C, ^15^N was produced as previously described (12). Briefly, transformed BL21 star (DE3) *E. coli* bacteria containing the appropriate vector codifying sfAFP were grown in 2L LB at 37°C until the OD600 reached 0.8 units. Afterwards, they were centrifuged and transferred to 0.5 L of minimal media with ^13^C-glucose and ^15^NH_4_Cl as the sole carbon and nitrogen sources, respectively. After one hour at 37 °C, expression was induced with 0.5 mM of IPTG and the culture was kept at 25 °C overnight. The protein was purified by Ni^++^ affinity chromatography followed by anionic exchange chromatography. Following purification, the His tag (sequence: MAHHHHHHVGTGSNDDDDKM) was not cleaved. The purified protein was then reduced with tris(2-carboxyethyl)phosphine (TCEP, obtained from Sigma), a strong, phosphine based reducing agent. Unlike β-mercaptoethanol, glutathione or dithiothreitol, TCEP does not form adducts with protein sulfhydral groups (18).

The NMR sample contained approximated 1.2 mM ^13^C, ^15^N sfAFP, 4.5 mM Na_2_HPO_4_/0.5 NaH_2_PO_4_, 3.0 mM TCEP, and 0.2 mM sodium trimethylsilylpropanesulfonate (DSS) as the internal chemical shift reference as well as 10% D2O for the spectrometer lock. Samples were placed in a water-matched Shigemi NMR tube and sealed with parafilm to reduce exposure to atmospheric oxygen.

The conformation and dynamics of reduced sfAFP were studied by NMR spectroscopy under three different sets of conditions, namely: 1) pH 2.5 & 25°C, 2) pH 2.5 & 5°C and 3) pH 6.15 & 5°C. The low pH conditions were chosen to optimize spectral quality for assignment, first at 25°C and then at 5°C, which is more physiologically relevant (13). The third set of conditions match those used to characterize oxidized, folded sfAFP (12).

### NMR Spectral Assignment

A Neo Avance 800 MHz (^1^H) NMR spectrometer, equipped with a ^1^H,^13^C,^15^N cryoprobe and Z-gradients, were utilized for most NMR experiments. The program Topspin (versions 2.1 and 4.0.8, Bruker Biospin) was utilized to record, transform and analyze the spectra. As sfAFP is disordered under the conditions employed here and since its sequence contains a high proportion of Gly and Ala residues, it is challenging for NMR spectral assignment. Since ^13^CO and ^15^N nuclei retain more dispersion in IDPs, we have used a non-conventional strategy based on a pair of 3D ^13^C-detected spectra which provide consecutive (*i, i+1*) ^13^CO and ^15^N backbone connectivities (19), (20). Even with this approach, some sequential residues with identical ^13^CO and ^15^N chemical shift values were observed. Therefore, another 3D ^1^H-detected spectrum which yields consecutive ^1^HN_*i*_-^1^HN_*i+1*_ connectivities was registered (21), (22). The analyses of all these data led to the essentially complete spectral assignment at pH 2.5 and 25°C. These results, and a ^1^H-^15^N HSQC spectrum recorded at 15°C, were used to transfer the assignments to pH 2.5, 5°C. Following the acquisition of spectra at pH 2.5, 5°C, the pH was increased to 6.15 by adding small volumes of 0.1 M Na2CO3 dissolved in D2O. A schematic diagram summarizing the assignment strategy is shown in **Sup. Fig. 2**. The initial analysis of the pH 6.15, 5°C data set revealed that the protein had been cleaved between residues T16/A17. This sample was also characterized to determine the effect of said cleavage on any residual structure and rigidity. Afterwards, another sample was prepared and the measurements at pH 6.15, 5°C were repeated.

#### Dynamic Measurements

{^1^H}-^15^N longitudinal relaxation rates (R_1_) and relaxation rates in the rotating frame (R_1ρ_) as well as the {^1^H}-^15^N heteronuclear NOE (hNOE) were recorded to assess dynamics on the ps/ns and μs/ms timescales. hNOE values measured for backbone amide groups as the ratio of resonance intensities registered in the presence or absence of saturation in an interleaved mode using a recycling delay of 11 seconds. Peaks were integrated with Topspin 4.0.8 (Bruker Biospin) and uncertainties were estimated as the standard deviation of the integrals of peak-sized spectral regions that lack signals and contain just noise.

Sets of seven ^1^H-^15^N correlation spectra with relaxation delays of 20, 1720, 1500, 860, 220, 1100 and 500 ms for R_1_ and ten ^1^H-^15^N correlation spectra with relaxation delays at 8, 300, 36, 76, 900, 100, 500, 156, 200 and 700 ms for R_1ρ_ were recorded. The R_1_ and R_1ρ_ relaxation rates and their estimated uncertainties were calculated using KaleidaGraph (version 3.6) by least-squares fitting of the exponential decay function I_t_ = I_0_·exp^(-kt)^, where It is the integral at time t, I0 is the integral at time zero, and k is the rate, to peak integral data measured with Topspin 4.0.8. In all cases, Pearson’s coorelation coefficient, R, was > 0.99.

## Results

### Reduced sfAFP is unfolded and lacks detectable populations of secondary structure

The 2D ^1^H-^15^N HSQC and ^13^C-^15^N CON spectra of ^13^C,^15^N-labeled sfAFP at pH 2.5 and 25 °C are shown in **Figure 1**. These spectra are strikingly different from those of oxidized, folded sfAFP observed at pH 6.15 and 5 °C (12). In particular, the excellent ^1^HN chemical shift dispersion observed for the folded protein is absent following reduction. Despite the low sequence complexity and poor ^1^HN chemical shift dispersion, by following the non-conventional assignment strategy described in the **Material and Methods** section, it was possible to obtain the essentially complete backbone ^1^HN, ^15^N, ^13^CO, ^13^Cα and side chain ^13^Cβ chemical shift assignments for reduced denatured sfAFP. The only unassigned resonances are ^1^H,^15^N for Cys1, ^13^Cα & ^13^Cβ for residues preceding proline residues (C13, G25, N40, TT70, A77, A80) and also ^13^Cα, ^13^Cβ, ^13^CO for P81. Assignments were also obtained at pH 2.5, 5°C (only backbone ^1^H-^15^N; 83% complete) and at pH 6.15, 5°C where 97% of ^1^HN, 97% of ^15^N, 96% of ^13^CO, 86% of ^13^Cβ and 98% of ^13^Cα signals are assigned. All three sets of assignments have been deposited in the BMRB databank under access number **51324**.

**Figure 1.**
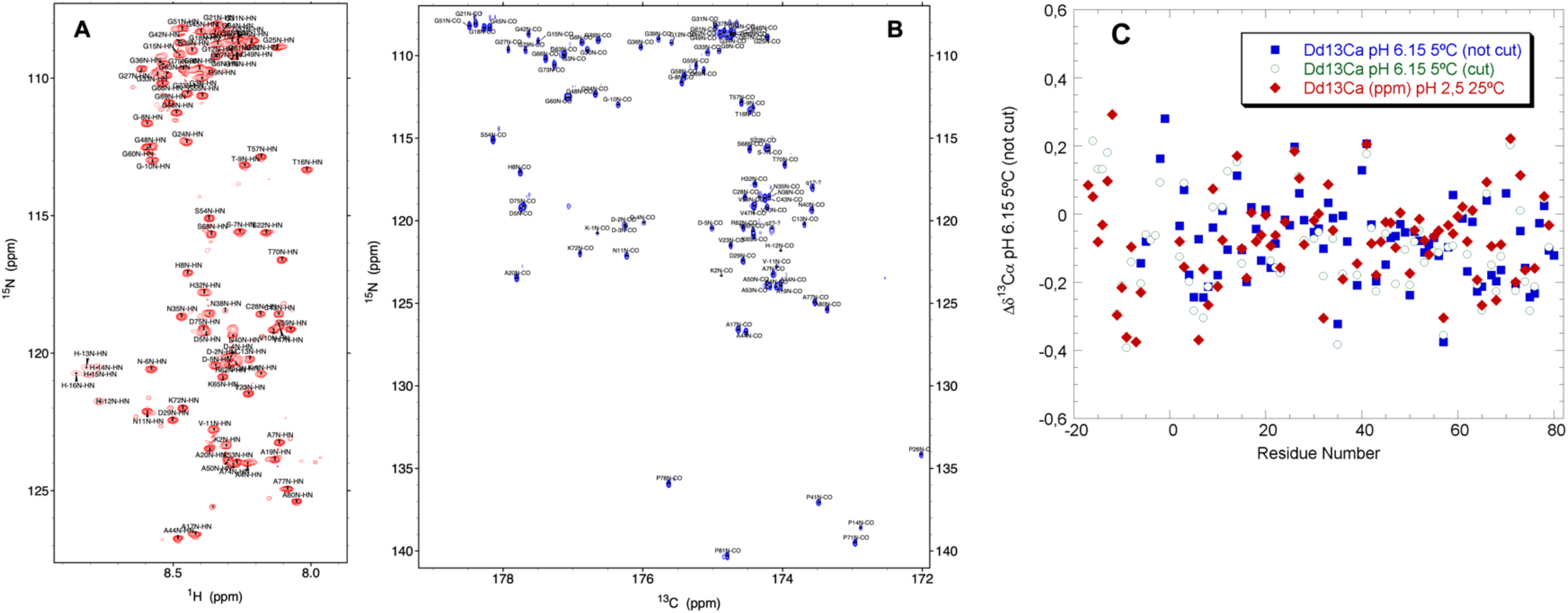
Assigned 2D ^1^H-^15^N HSQC and 2D CON spectra and ^13^Cα conformational chemical shifts show that reduced ^13^C,^15^N-sfAFP is thoroughly denatured. Assigned 2D ^1^H-^15^N HSQC (**red**, **A**) and 2D ^13^CO^15^N (**blue**, **B**) of reduced ^13^C,^15^N sfAFP at pH 2.5, 25°C. The ^15^N of each residue are labeled. Residues of the His tag are labeled with negative numbers. **C**. Conformational ^13^Cα chemical shifts for sfAFP at pH 2.5, 25°C (red), and at pH 6.15 and 5°C with (**green**) or without (**blue**) a proteolytic cleavage between residues T16 / A17.

The observation of signals of reduced TCEP in 1D, ^1^H spectra (**Sup. Fig. 3**) means that the Cys will be reduced since TCEP is a potent reducing agent and quantitatively reduces disulfide bonds (18). The ^13^Cβ chemical shift values of the Cys residues are observed to be < 30 ppm. Taking into account reported ^13^Cβ chemical shift values for reduced (< 32 ppm) and oxidized (> 35 ppm) Cys (23), these values indicate that the four Cys side chains are reduced to -CH2-SH and that the two disulfide bonds are broken. sfAFP contains six proline residues, four of which paradoxically lie in turns outside the six PPII helices. Whereas two of the Pro residues adopt the *cis* conformation in folded sfAFP, here all ^13^Cβ proline chemical shifts range from 32.0 – 32.3 ppm and are consistent with the *trans* conformation (24). This concords with unfolding as the *trans* conformation normally dominates denatured state ensembles. Following the example of (25), we analyzed minor peaks but none arising from Xaa-Pro *cis* peptide bonds could be rigorously assigned.

Reduced, denatured sfAFP might retain some partially structured segments as has often been observed in intrinsically disordered (26) or chemically denatured proteins (15) (27). Since ^1^HN chemical shift dispersion arises mainly from >C=Oooo^1^H-N H-bond formation, the loss of dispersion observed for all the conditions studied here indicates that the interhelical H-bond network has broken down following reduction. To further test for residual structure, the conformational chemical shifts (Δδ) for ^13^Cα plotted are in **Fig. 1C** for reduced sfAFP at three conditions; *i.e*. i) pH 2.5 & 25°C; ii) pH 6.15 & 5°C for the sample containing a proteolytic cleavage between T16 and A17, or iii) pH 6.15 & 5°C without that cleavage. The mean Δδ ^13^Cα values are close to zero for all three data sets; namely: −0.07 ± 0.14; −0.08, ± 0.12; and −0.09 ± 0.15 ppm, respectively. Moreover, there is no statistical difference between the three sets of Δδ^13^Cα values as judged by p-values > 0.40 for each pairwise comparison. Based on these findings, we conclude that reduced, denatured sfAFP is essentially completely unfolded following reduction.

#### sfAFP retains some rigidity on ps/ns timescales following reduction

To assess the ps/ns flexibility of reduced denatured sfAFP, we measured the {^1^H}-^15^N NOE ratios for residues whose ^1^H-^15^N signals are well resolved in the 2D ^1^H-^15^N HSQC spectra. Whereas these ratios are high in the folded protein and even approach values indicative of complete rigidity (12), the values in the reduced, denatured protein are much lower (**Fig. 2 ABC**). Overall they are about halfway between values expected for fully stiff (0.85) and high flexibility. Similar {^1^H}-^15^N NOE ratios were observed in three different conditions: pH 2.5, 25°C; pH 2.5, 5°C and pH 6.15, 5°C. The N-terminal His tag also appears to be somewhat more rigid than the rest of denatured sfAFP and the residues corresponding to the last PPII helix seem to be more flexible. Within the rest of the protein, *i.e*. residues 1-70, there seem to be a slight trend towards increased flexibility moving from the N-terminus to the C-terminus. This trend might be related to a lower content of potentially stabilizing charge-charge interactions in the C-terminal half of sfAFP.

**Figure 2:**
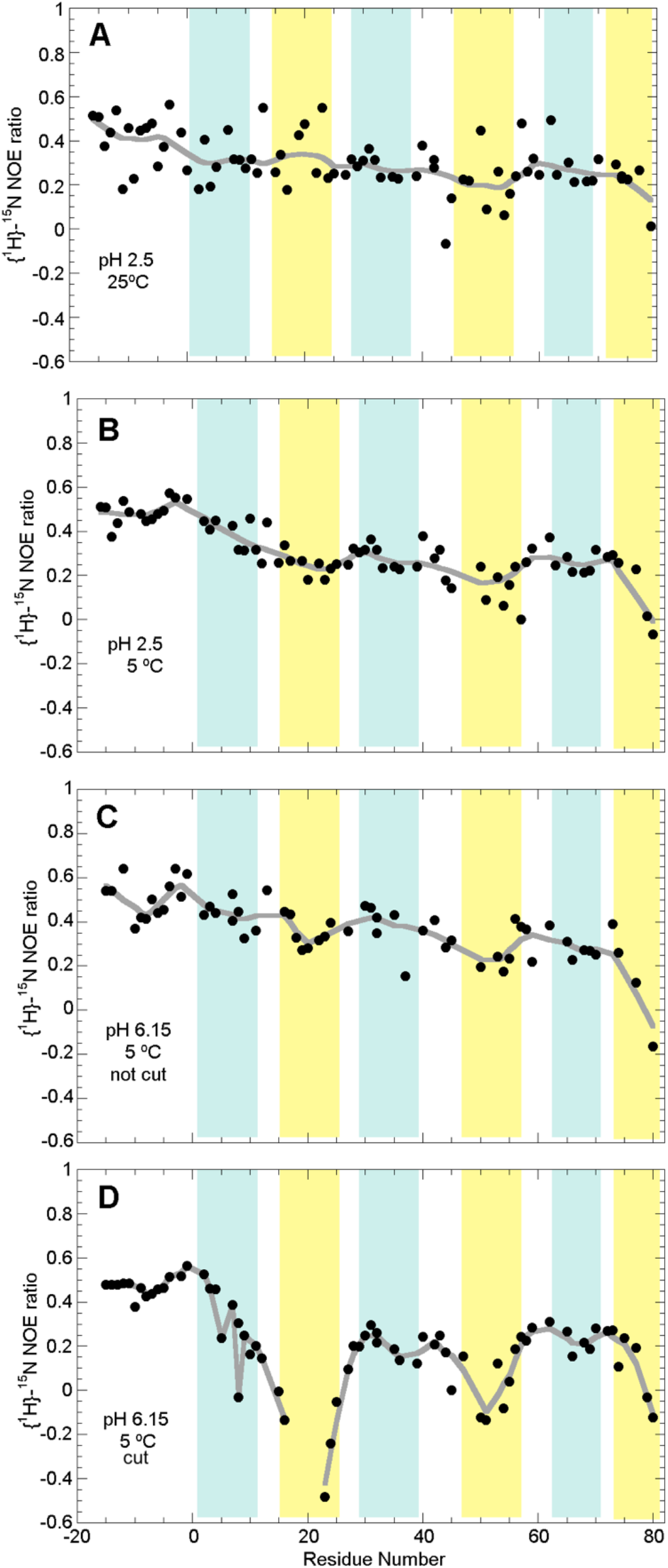
Fast ps/ns dynamics of reduced denatured sfAFP. The per-residue {^1^H}-^15^N ratios are shown for reduced, denatured sfAFP at: **A**. pH 2.5, 25°C, **B**. pH 2.5, 5°C, **C**. pH 6.15, 5°C and **D**. pH 6.15, 5°C with an internal proteolytic cleavage. Shaded regions show the approximate positions of polar (cyan) and nonpolar (yellow) in PPII helices in the folded protein. The gray line marks the trend as calculated with a weighting function.

In the presence of the proteolytic cleavage, a dramatic decrease in the {^1^H}-^15^N NOE values is seen in the nearby residues corresponding to the second PPII helix, *i.e*. the helix which contains the cleavage site, and a moderate decrease is also seen in the residues corresponding to the fourth PPII helix (**Fig. 2 D**). Considering that both PPII helices 2 and 4 harbor nonpolar residues, these results suggests that hydrophobic contacts in the denatured state might reduce the flexibility of the uncleaved, reduced denatured sfAFP. Similar hydrophobic clusters in denatured proteins been reported previously (27).

#### Rigidity on μs/ms timescales increases upon cooling and is highest in the N-terminal polar segments

The longitudinal relaxation rates (R_1_) and relaxation rates in the rotating frame (R_1ρ_) were measured for reduced, denatured sfAFP under three different conditions: pH 2.5, 25°C; pH 2.5, 5°C; and pH 6.15, 5°C to assess mobility on the μs/ms timescales and the results are shown in **Figure 3**. Overall, the rates are low, which is in line with the denatured character of the chain under reducing conditions. Some interesting trends can be observed. First, the residues of the N-terminal His-tag show relatively high R_1ρ_ rates, which may be due to a dearth of Gly residues. Secondly, the mobility, gauged most clearly by R_1ρ_ values, diminishes as the temperature is lowered from 25°C to 5°C (**Figure 3 A** versus **B**) and when the pH is increased from 2.5 to 6.15 (**Figure 3**, **B** versus **C**). This increase is especially noticeable for the residues that correspond to PPII helices 1 and 3 in the folded protein. This suggests that the balance of electrostatic interactions may favor certain conformations at neutral pH, stiffening the chain. Thirdly, under all three conditions, the residues corresponding to the first PPII helix have higher R_1ρ_ rates, meaning they are more rigid, whereas the rates for the C-terminal residues are lower. Moreover, the residues which would constitute the polar PPII helices in the folded protein (**Sup. Fig. 1, Fig. 3**), tend to be more rigid than those corresponding to the nonpolar PPII helices. The results obtained at pH 6.15, 5°C, which are approximately the physiological conditions that the nascent sfAFP chain would experience after synthesis, is suggestive of some innate rigidity in the residues corresponding to PPII helices 1 and 3 which may help guide nascent folding.

**Figure 3:**
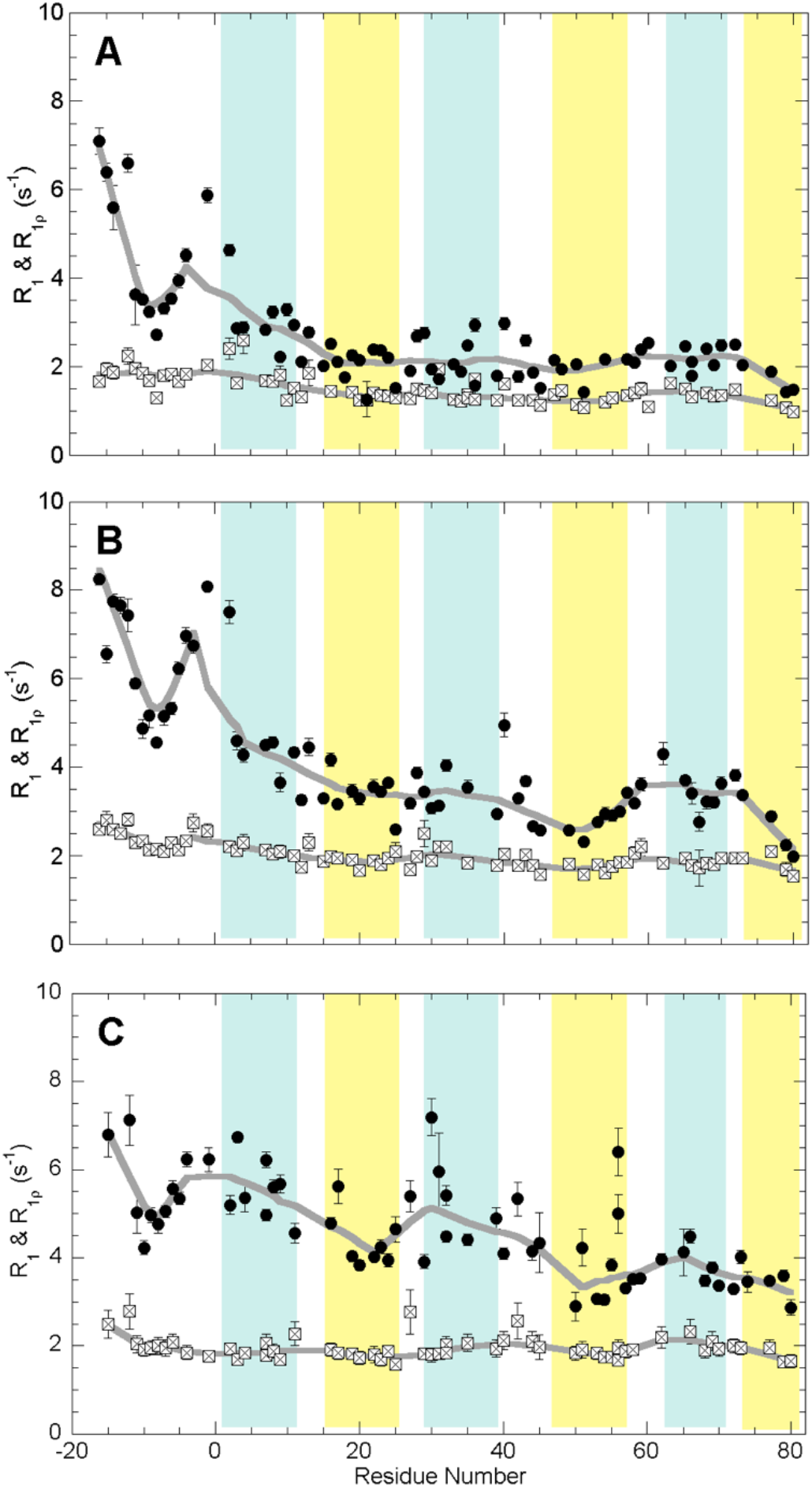
R_1_ and R_1ρ_ relaxation rates of reduced denatured sfAFP. The per-residue {^1^H}-^15^N longitudinal relaxation rates (R_1_, open squares) and longitudinal relaxation rates in the rotating frame (R_1ρ_, filled circles) are shown for reduced, denatured sfAFP at: **A**. pH 2.5, 25°C, **B**. pH 2.5, 5°C and **C**. pH 6.15, 5°C. Shaded regions show the approximate positions of polar (cyan) and nonpolar (yellow) in PPII helices in the folded protein. The gray line marks the trend as calculated with a weighting function.

## Discussion

Research on the sfAFP is adding valuable understanding to the folding and conformational stability of proteins composed by polyproline II helical bundles. Our study builds on investigation from the Sosnick lab, who used folding kinetics, SAXS and computational modeling of disulfide-bond intact (*i.e*. -S—S-) sfAFP to shed light on the energetics and folding mechanism of sfAFP (13), and on our NMR and computational energetics study of disulfide-bond intact, folded sfAFP (12) which elucidated the native state dynamics and additional stabilizing contributions. These two studies drew a few different conclusions; namely, whereas Gates *et al*. (13) reported that conformational entropy of sfAFP’s folded state is higher than that of typical proteins composed of α-helices or β-sheets, Treviño *et al*, (12) reported NMR relaxation measurements suggesting that folded sfAFP is just as stiff as other folded proteins. Moreover, while Gates *et al*. (13) reasoned that the stabilizing contribution of the hydrophobic effect is low, Treviño *et al*. (12) uncovered that sfAFP buries some nonpolar surface area through dimerization. On the other hand, and more importantly, both studies concurred that Cα-HooO=C H-bonds also contribute to native state stability. Gates *et al*., found that the high Gly content does not make the unfolded ensemble of sfAFP especially condensed but that this ensemble is biased towards PPII conformation (13). This conformational bias might be related to the moderate stiffness found here for the reduced, denatured state on ps/ns timescales. Finally, Gates *et al*., reasoned that the conformational entropy change during folding (DS_conf·F_) is higher for glycine residues due to its more flexible backbone. However, since glycine residues do not have a side chain, they pointed out that this decreases the ΔS_conf·F_ (13). The sum of these two effects is that glycine residues’ contribution to the ΔS_conf·F_ is similar to that of non-Gly residues.

Our results, obtained under physiologically relevant reducing conditions corroborate the model for sfAFP folding with disulfide bonds present proposed by Gates, Sosnick and co-workers (13) (**Figure 4**). As they advanced, proline residues could favor turn forming, preorganizing the chain into segments corresponding roughly to the PPII helices. Whereas aspartic acid residues are neutral at pH 3, they are predominantly charged at pH 6 where our relaxation measurements found increased rigidity. These findings lead us to additionally suggest that electrostatic interactions formed by K2, D5, H8, D29 and H32 are present at pH 6 and might aid the juxtaposition of C1 and C28, favoring disulfide bond formation. This would enhance the folding of PPII helices 1 and 3, and may position C13 and facilitate its disulfide bond formation with C43. Proline *cis/trans* isomerization impacts protein folding and function(15) (28). Here, the formation of this second disulfide bond might be rate limited by the isomerization of the C13-P14 peptide bond from *trans*, its predominant conformation in the unfolded state, to the *cis* conformation observed in the native state. By contrast, the folding of the remaining polar PPII helix (helix 5), may be disfavored by repulsive electrostatic interactions among H32, R62 and K65. Whereas the both the first and the last PPII helices only interact with two other PPII helices each, only the first is stabilized by two disulfide bridges. The presence of a potential stabilizing charge – charge interaction between K72 and D75 and two Pro residues P78 and P81 could stabilize the last PPII helix. This is supported by results on the larger 203-residue-long isoform of the snow flea antifreeze protein which is modeled to be composed of thirteen PPII helices (29). This longer isoform conserves the first disulfide bond (C1-C26) and two prolines in the twelfth PPII helix.

**Figure 4:**
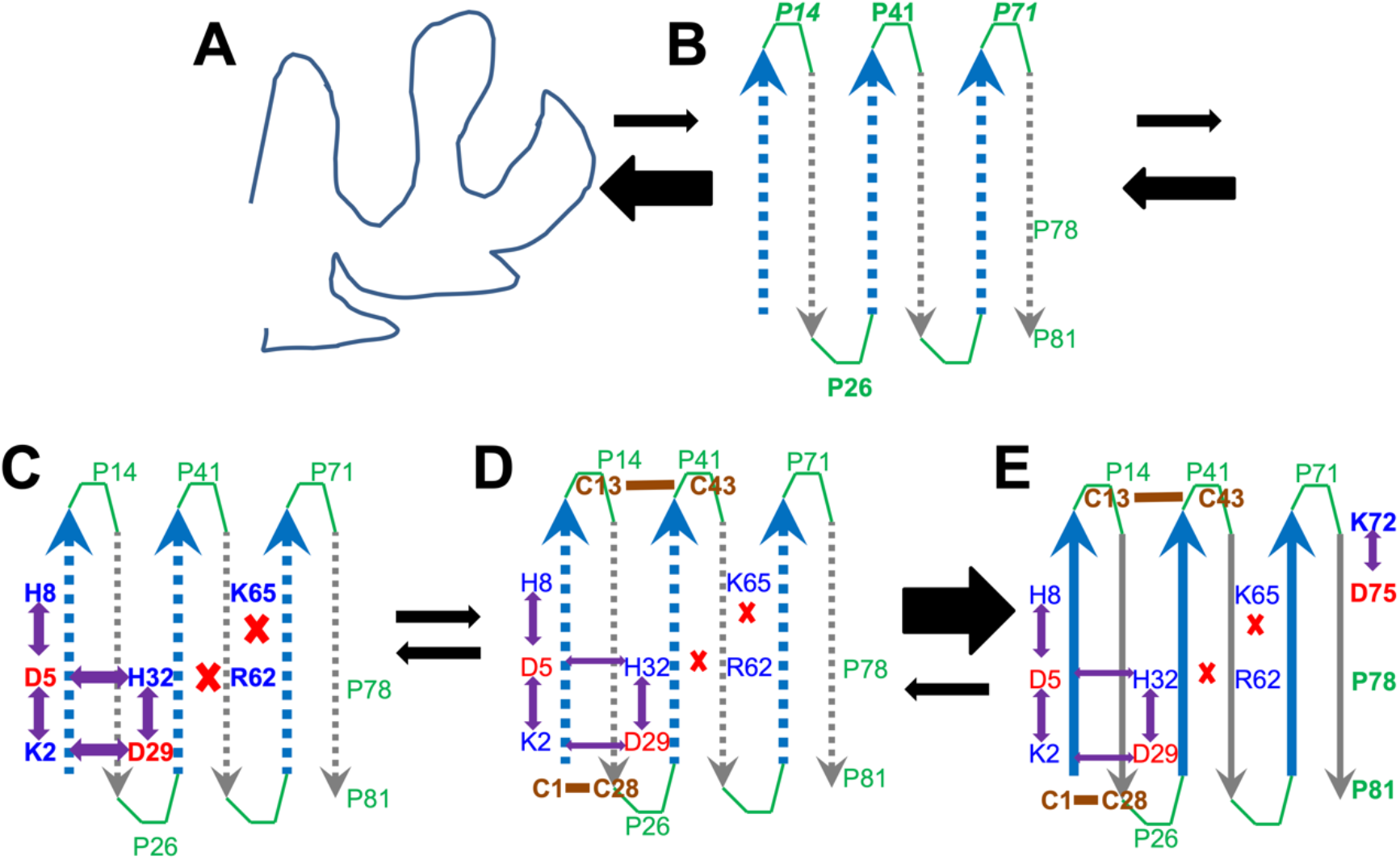
Events in sfAFP Folding Inferred from Dynamics. Within the disordered ensemble (**A**), proline residues *14*, 26, 41 and *71* predispose the chain to form turn-like conformers, though prolines *14* and *71*, which adopt *cis* Xaa-Pro peptide bonds in native sfAFP, may slow folding (**B**). A small network of favorable charge-charge interactions, inferred from higher R_1ρ_ rates at pH 6.15 versus 2.5, which would form among residues that will constitute PPII helices 1 and 3 (blue dottted arrows) (**C**) might help colocalize the Cys1 and Cys28 to aid disulfide bond formation; this could then help position C13 close to C43 (**D**). The lack of stabilizing charge/charge interactions in the helix containing R62 and K65 is consistent with its higher dynamics. Disulfide bond formation decisively stabilizes the native state consisting of three polar PPII helices (blue) and three nonpolar PPII helices (gray). The sixth PPII (**E**), which forms fewer interhelical contacts, is stabilized locally by the presence of a charge-charge interaction between Lys 72 and Asp 75 and two additional Proline residues (78 and 81). Further stabilization would be afforded by hydrophobic contacts upon dimerization (12); this is not represented in this diagram.

Here, we found a moderate stiffness seen in reduced, denatured sfAFP, which strongly suggests that the unfavorable ΔSconfoF will be lower than if the protein were more flexible under denaturing conditions. It would be interesting to precisely measure this ΔS_conf·F_ for sfAFP and compare it to values for typical proteins with an average glycine residue content and composed of α-helices and/or β-sheets. However, a quantitative calculation for ΔS_conf·F_ based on NMR relaxation data faces important limitations, as previously discussed (30)(31). Nevertheless, the hNOE, R_1_ and R_1ρ_ values measured here are similar to those reported previously under denaturing conditions for representative proteins such as hen egg white lysozyme (14), CheY (32) and *Staphylococcal* nuclease (33) or protein domains like an SH3 domain (34). Taken together with the similar rigidities of the folded states of typical glycine-poor, α, β, α+β proteins and sfAFP (12), this suggests that the ΔS_conf·F_ of all these proteins would be similar despite the high glycine content of the latter. This means that the free energy “cost” paid by glycine-rich PPII helical bundle proteins for losing ΔS_conf·F_ is likely similar to that paid by typical proteins. Therefore, glycine-rich PPII helical bundle proteins should not require an exceptionally robust set of stabilizing interactions to fold and be stable. A “normal” set typical of an “average” α/β protein should do. This finding should facilitate the *de novo* design of PPII helical bundles.

In conclusion, our results show that reduced sfAFP is fully denatured upon reduction of its two disulfide bonds. Despite a high proportion of glycine residues, its denatured ensemble is not more flexible than those of typical proteins composed of α-helices and/or β-sheets. The insight will help guide the design of new proteins composed of polyproline II helical bundles.

---

**Sup. Fig. 1:**
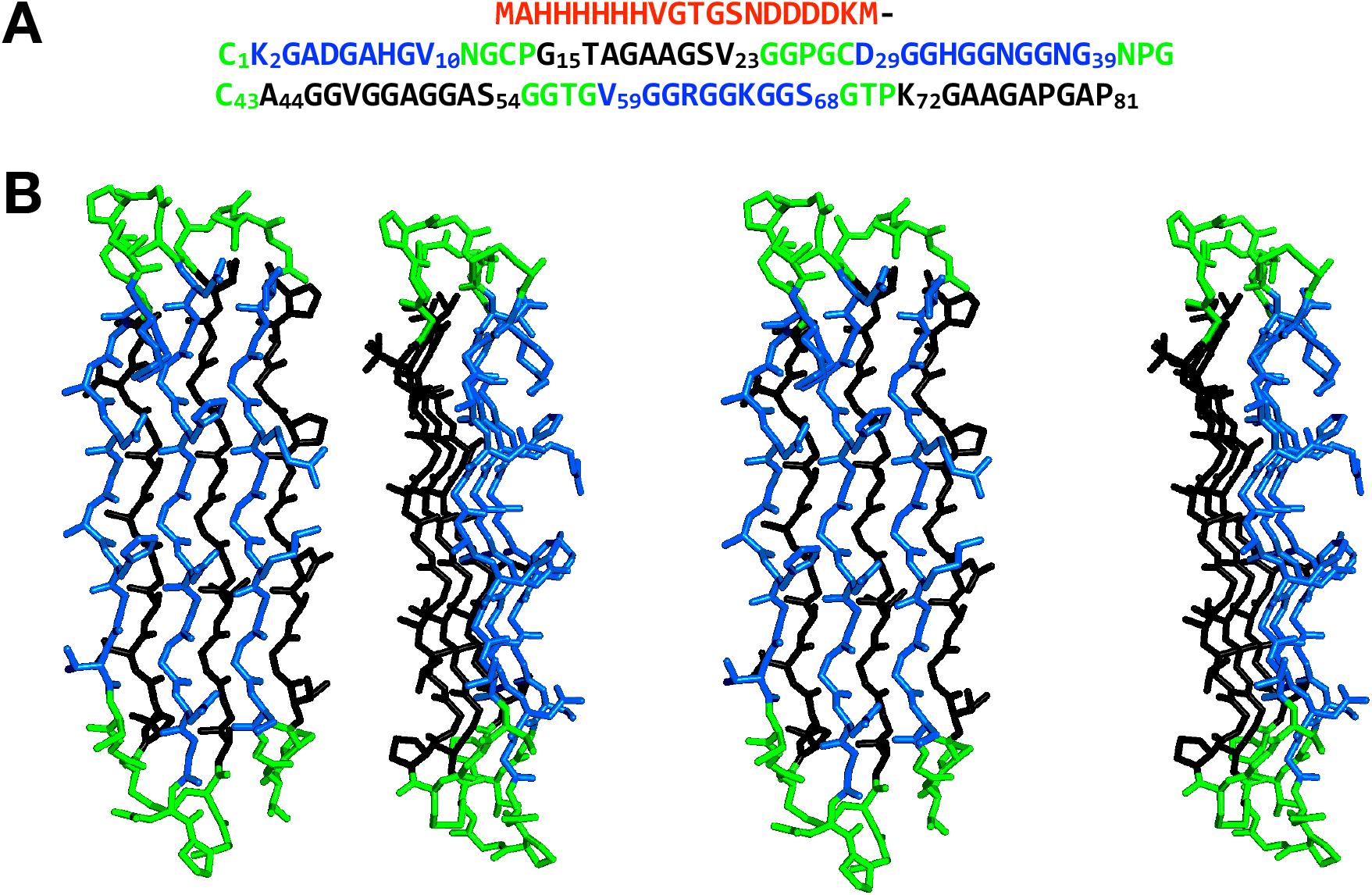
Sequence and 3D structure of sfAFP. **A.** Sequence of sfAFP. The His-tag is colored red. Residues belonging to the loops are colored green, those forming PPII helices on the polar face are colored blue and PPII helices on the hydrophobic face are colored black. **B.** Crosseyed stereo diagrams of the sfAFP structure (two perspectives) with the residues colored as in panel **A.**

**Sup Fig. 2.**
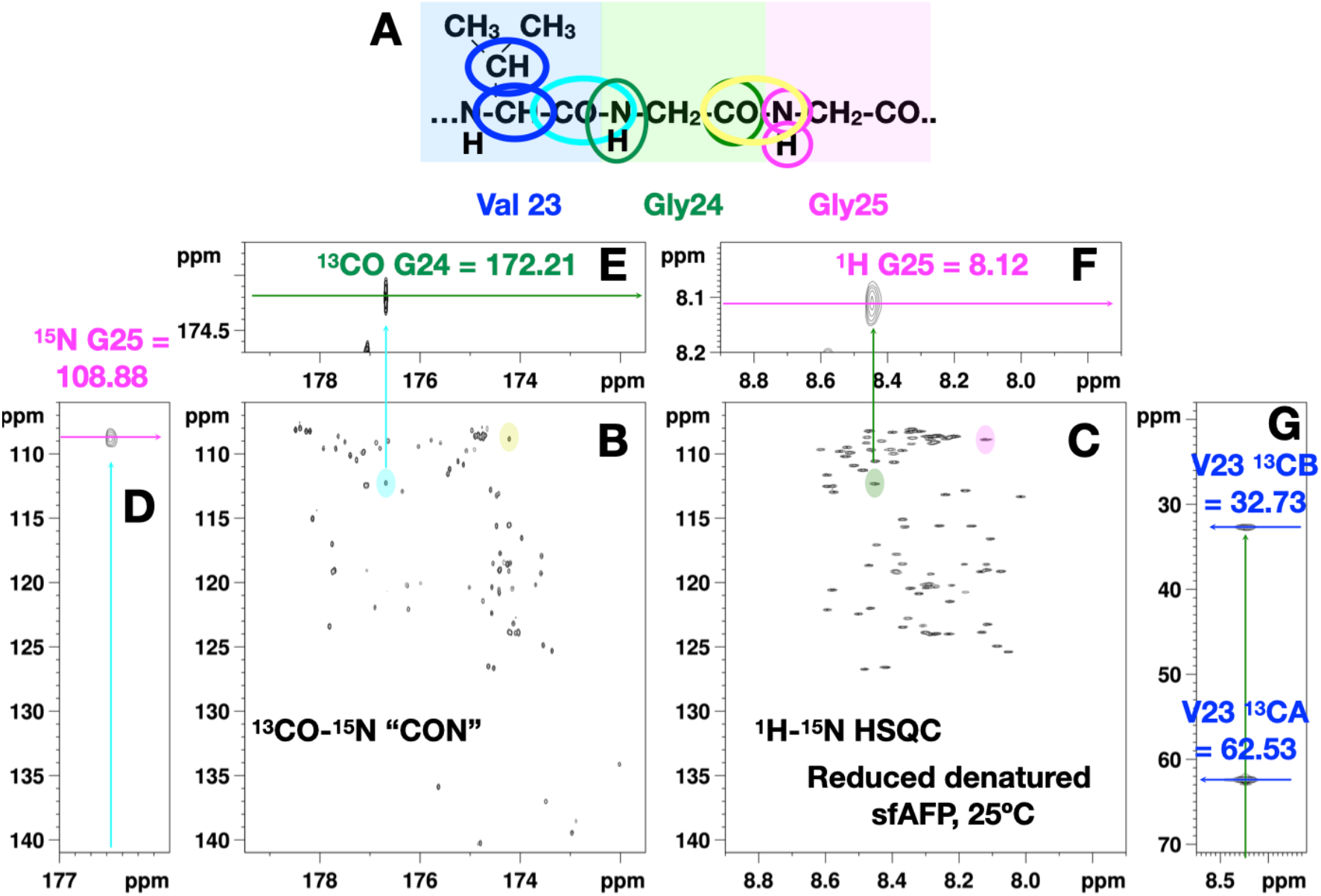
Sequential Assignment Strategy for Reduced sfAFP. To assign a segment of reduced ^13^C,^15^N-sfAFP (panel **A**), we start with an isolated peak (shaded **cyan**) in the 2D ^13^CO-^15^N spectrum (panel **B**). Unless it arises from a Proline residue, each CON peak will have a sister signal (shaded **green**) in the 2D ^1^H-^15^N HSQC (panel **C**) which shares the same ^15^N δ value. Using the ^13^CO and ^15^N δ values of the starting peak, the δ values of the next ^15^N and ^13^CO nuclei along the chain are identified in two 3D hacocoNcaNCO and hacaCOncaNCO spectra, whose strips are shown in paneles **D** and **E**, respectively. This permits the next ^13^CO^15^N correlation (panel **B**, shaded yellow) in the 2D CON spectra to be identified. Likewise, the δ value of the next ^1^HN nuclei to the sister ^1^H-^15^N peak is identified in a 3D HncocaNH spectrum (panel **F**), affording the next ^1^H-^15^N signal along the segment (panel **C**, shaded **lilac**). Finally in a 3D CBCACON spectrum (panel **G**), the resonances of the ^13^Cα and ^13^Cβ nuclei of the first residue are found and corroborate the assignment. Once determined at pH 2.5, 25°C, the transfer of the assignments 5°C was aided by recording an ^1^H-^15^N HSQC spectrum at 15°C. Because some peaks move in a nonlinear fashion when the conditions were changed to pH 6.15, 5°C, 2D ^1^H-^15^N HSQC, 2C CON as well as 3D HNCO and CBCAcoNH spectra were recorded and analyzed to determine the assignments.

**Sup. Fig. 3.**
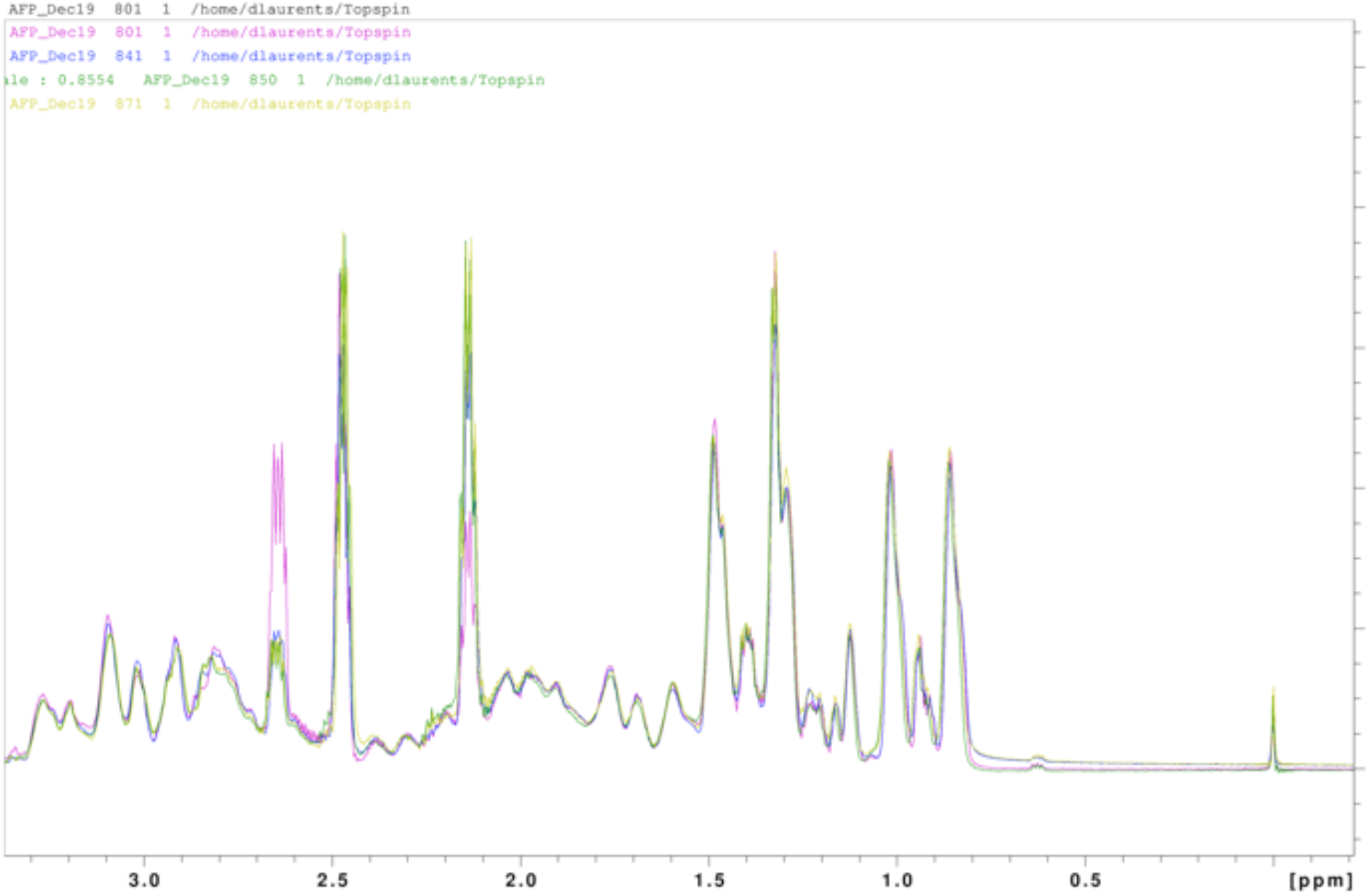
Observation of Signal from Reduced TCEP. Four 1D ^1^H NMR spectra (colored purple, blue, green and amber from first to last) of ^13^C,^15^N reduced denaturated sfAFP at pH 2.5, 5°C recorded successively over the course of the 2D and 3D spectral adquisition. The earliest spectrum (purple) shows an intense peak at 2.64 ppm which corresponds to reduced TCEP. Over time, this peak weakens and a peak corresponding to oxidized TCEP (2.13 ppm) strengthens. Nevertheless, the peak at 2.64 ppm indicating that some reduced TCEP is always present, ensuring that the Cys remain reduced. The peak at 2.47 ppm contains a combination of reduced and oxidized signals and the signals at 0.00 (intense singlet) and 0.63 (weak triplet) correspond to the trimethyl and a methylene groups, respectively of sodium trimethylsilylpropanesulfonate (DSS) used for chemical shift referencing.

